# Introducing ribosomal tandem repeat barcoding for fungi

**DOI:** 10.1101/310540

**Authors:** Christian Wurzbacher, Ellen Larsson, Johan Bengtsson-Palme, Silke Van den Wyngaert, Sten Svantesson, Erik Kristiansson, Maiko Kagami, R. Henrik Nilsson

## Abstract

Sequence analysis of the various ribosomal genetic markers is the dominant molecular method for identification and description of fungi. However, there is little agreement on what ribosomal markers should be used, and research groups utilize different markers depending on what fungal groups are targeted. New environmental fungal lineages known only from DNA data reveal significant gaps in the coverage of the fungal kingdom both in terms of taxonomy and marker coverage in the reference sequence databases. In order to integrate references covering all of the ribosomal markers, we present three sets of general primers that allow the amplification of the complete ribosomal operon from the ribosomal tandem repeats. The primers cover all ribosomal markers (ETS, SSU, ITS1, 5.8S, ITS2, LSU, and IGS) from the 5’ end of the ribosomal operon all the way to the 3’ end. We coupled these primers successfully with third generation sequencing (PacBio and Nanopore sequencing) to showcase our approach on authentic fungal herbarium specimens. In particular, we were able to generate high-quality reference data with Nanopore sequencing in a high-throughput manner, showing that the generation of reference data can be achieved on a regular desktop computer without the need for a large-scale sequencing facility. The quality of the Nanopore generated sequences was 99.85 %, which is comparable with the 99.78 % accuracy described for Sanger sequencing. With this work, we hope to stimulate the generation of a new comprehensive standard of ribosomal reference data with the ultimate aim to close the huge gaps in our reference datasets.

## Introduction

In 1990 it became clear that ribosomes are common to all extant organisms known today (Woese et al. 1990). The ribosomal genetic markers are located in the ribosomal operon, a multi copy region featuring both genes, spacers, and other poorly understood elements (Rosenblad et al. 2016). Clearly defined fragments of these regions and spacers have been identified as suitable markers for various scientific pursuits in different groups of organisms (Hillis and Dixon 1991; Tedersoo et al. 2015), due to their individual substitution rates and length. Well-known examples include the SSU, which has been used to explore the phylogeny of prokaryotes and microeukaryotes, and the ITS region, which is the formal barcode for molecular identification of fungi (Schoch et al. 2012).

The fungal kingdom is vast, with estimates ranging from 1.5 to 6 million extant species (Taylor et al. 2015; Hawksworth and Lücking 2017). On the other hand, a modest 140,000 species have been described so far (http://www.speciesfungorum.org/Names/Names.asp, accessed March 2018), underlining a considerable knowledge gap. From environmental sequencing, the discrepancy between the known and unknown fungi becomes readily apparent, where it is not uncommon to find for instance that >10% of all fungal species hypotheses (a molecular based species concept akin to operational taxonomic units, OTUs; Blaxter et al. 2005; Kõljalg et al. 2013) do not fall in any known fungal phylum (Nilsson et al. 2016). This hints at the presence of a large number of unknown branches on the fungal tree of life (Tedersoo et al. 2017a). Compounding this problem, not all described fungi have a nucleotide record, which is often but not always related to older species descriptions made before the advent of molecular biology (Hibbett et al. 2016). Consequently, many studies fail to classify more than 15-20% of the fungal sequences to genus level (e.g., Wurzbacher et al. 2016), which severely hampers the interpretation of these results.

While the ITS is a well chosen barcode, it is less suitable for phylogenetic analysis and it is not optimal for all fungal lineages. For instance, the Cryptomycota taxonomy is based on the ribosomal SSU as the primary genetic marker (e.g. Lazarus et al. 2015), while Chytridiomycota taxonomists mainly work with ribosomal large subunit data (LSU) (Letcher et al. 2006). Similarly, studies on Zygomycota often employ the SSU and LSU (White et al. 2006), while work on yeast species is often done using the LSU (Burgaud et al. 2016). Researchers studying fungal species complexes regularly need to consider genetic markers with even higher substitution rates than the ITS (e.g., the IGS region, O’Donnell et al. 2009; Nilsson et al. 2018).

There are thus serious gaps in the reference databases relating to 1) taxonomic coverage and 2) marker coverage. Some groups have ample SSU data; others have a reasonable ITS and LSU coverage; others are known only from ITS or LSU data. We argue that it is crucial to close these two types of gaps - ideally at the same time – to achieve a robust data-driven progress in mycology. Having access to all ribosomal markers at once solves a range of pertinent research questions, such as trying to obtain a robust phylogenetic placement for an ITS sequence (e.g. James et al. 2006), trying to prove that an unknown taxon does indeed belong to the true fungi (Tedersoo et al. 2017), or moving forward in spite of the fact that an ITS sequence produces no BLAST matches at all in INSDC databases (Heeger et al. 2018). Thus, an urgent goal is to fill the taxonomic and marker-related gaps in our reference sequence databases (e.g. SILVA: Glöckner et al. 2017, UNITE: Kõljalg et al. 2013, and RDP: Cole et al. 2014).

In this study we explore one promising way to close the gaps in the reference databases and simultaneously unite them through the use of the emerging long-read sequencing technologies. Here, we aim to generate high-quality *de novo* reference data for the full ribosomal operon and the adjunct intergenic regions, which would unify five or more distinct marker regions: the SSU gene, the ITS including the 5.8S rRNA gene, the LSU gene, and the IGS (that often contains the 5S gene, too). Fortunately, the eukaryotic ribosomal operon is arranged in tandem repeats in the nuclear genome, which makes its amplification by PCR comparatively straightforward. In theory at least, DNA sequencing of the full ribosomal operon is perfectly possible. So far, it has not been feasible to sequence such long DNA stretches in a simple, time and cost-efficient way. In principle, Sanger sequencing with maybe 10 internal sequencing primers is a possibility, or alternatively shot-gun sequencing, which requires prior fragmentation of the long amplicon. Regardless, the latter methods are less than straightforward by requiring substantial time, multiple rounds of sequencing, and significant laboratory expertise.

In contrast, emerging third-generation sequencing technologies - MinION (Oxford Nanopore Technologies; https://nanoporetech.com/) and PacBio SMRT sequencing (Pacific Biosciences; http://www.pacb.com/) - offer the possibility to sequence long DNA amplicons in a single read, very much like Sanger does so well for short amplicons. Both of these technologies are suitable for high-accuracy, long-range sequencing (Singer et al. 2016; Benitez-Paez and Sanz 2017; Tedersoo et al. 2018; Karst et al. 2018). PacBio excels where high accuracy is needed due to its circular consensus sequencing mode. The advantage of Nanopore sequencing is the price, and the fast processing time. In addition, there is no need for a sequencing provider, since the sequencing can be done at a regular desktop computer. We think that these features combine to make both technologies invaluable for our envisioned generation of comprehensive reference data.

In this work, we present PCR primers to cover the whole fungal nuclear ribosomal region in either two shorter amplifications of 5 kb each or in a single long amplicon of approximately 10 kb. The end product of both approaches is a 10 kb long stretch of nucleotide data that comprises all ribosomal markers, thus forming reference sequence data that satisfy many different research questions at once. Our secondary objective is to provide a cost-efficient and easy-to-use system that can be adopted even in small laboratories with limited budgets to facilitate broad generation of complete ribosomal reference data that may, as a joint effort, eventually help to fill our knowledge gaps in mycology and elsewhere.

## Methods

### Tested samples

We tested several samples of various origins to evaluate the primer systems for our respective fields of research with a focus on reference material from herbarium samples (Supplemental Material S1). For Basidiomycota species within our target class of Agaricomycetes, we tested DNA extracted from herbarium specimens deposited at the infrastructure of University of Gothenburg, Herbarium GB (n = 66). DNA extraction from fungal material was performed with the DNA Plant Mini Kit (Qiagen). We furthermore evaluated the use of the primers for a few early diverging, poorly described, environmental fungal lineages. For parasitic uncultured aquatic fungi (Chytridiomycota), we employed micromanipulation as described in Ishida et al. (2015). This was followed by a whole genome amplification (illustra single cell GenomiPhi V1/V2 DNA amplification kit; GE Healthcare), which served as DNA template for our ribosomal PCRs (n = 9). Furthermore, to test the amplification of extremely distant fungal lineages from an animal host, we worked with DNA extracted from Malpighian tubule tissue from two cockroaches infected with *Nephridiophaga,* derived from previous work (Radek et al. 2017).

### Primer design

The operon PCR was performed with newly designed and adapted primer pairs. The primer pairs were modified from previous primers, namely the universal NS1 primer that offers a broad coverage of many eukaryotic lineages (White et al. 1990) and a Holomycota/Nucletmycea-specific primer derived from the RD78 primer (Wurzbacher et al. 2014). The modified and further developed primers were designed and tested in ARB v. 6 (Ludwig et al. 2004) against the SILVA reference databases v. 123 (Glöckner et al. 2017) for SSU and LSU. NS1 was shortened by three bases at the 3’ end to avoid mismatches with major Chytridiomycota and Cryptomycota lineages. Furthermore, to maintain an acceptable melting temperature, the primer was prolonged with two bases at the 5’ end from a longer version of NS1 mentioned in Mitchell & Zuccharo (2011), and was named NS1short: CAGTAGTCATATGCTTGTC^1^. We further developed the RD78 primer with the Probedesign and Probematch tools integrated in ARB, by shifting it towards the LSU 5’ by several bases, so that the mismatches to outgroups such as Eumetazoa fall in the 3’ end of the primer region. This will facilitate the application of the primer in, e.g. mixed template samples such as environmentally derived DNA. Similar to RD78, the resulting primer is highly specific to true fungi, and we named it RCA95m (CTATGTTTTAATTAGACAGTCAG), since it covers more than 95% of all fungi in the SILVA LSU database. However, we found exceptions (mismatches) in a few long-branched Zygomycota (e.g., *Dimargaris)* and in the close vicinity to the genus *Neurospora* (Ascomycota). This primer pair (NS1short and RCA95m) was used to amplify the greater part of the ribosomal operon (rDNA PCR, see Figure 1) of the ribosomal tandem repeat. In order to amplify the missing parts of the ribosomal region (the 5’ end of the LSU, IGS, and the ETS region), we simply used the reverse complementary version of each primer: NS1rc (ACAAGCATATGACTACTG) and RCA95rc (CTGACTGTCTAATTAAAACATAG). As a third primer pair we developed a primer pair based on RCA95m that binds to a single position in the LSU; the forward primer Fun-rOP-F (CTGACTGTCTAATTAAAACAT) amplifies in the 3’ direction of the LSU, while the reverse primer Fun-rOP-R (TCAGATTCCCCTTGTCCGTA) amplifies in the 5’ direction (see Figure 1).

**Figure 1.**
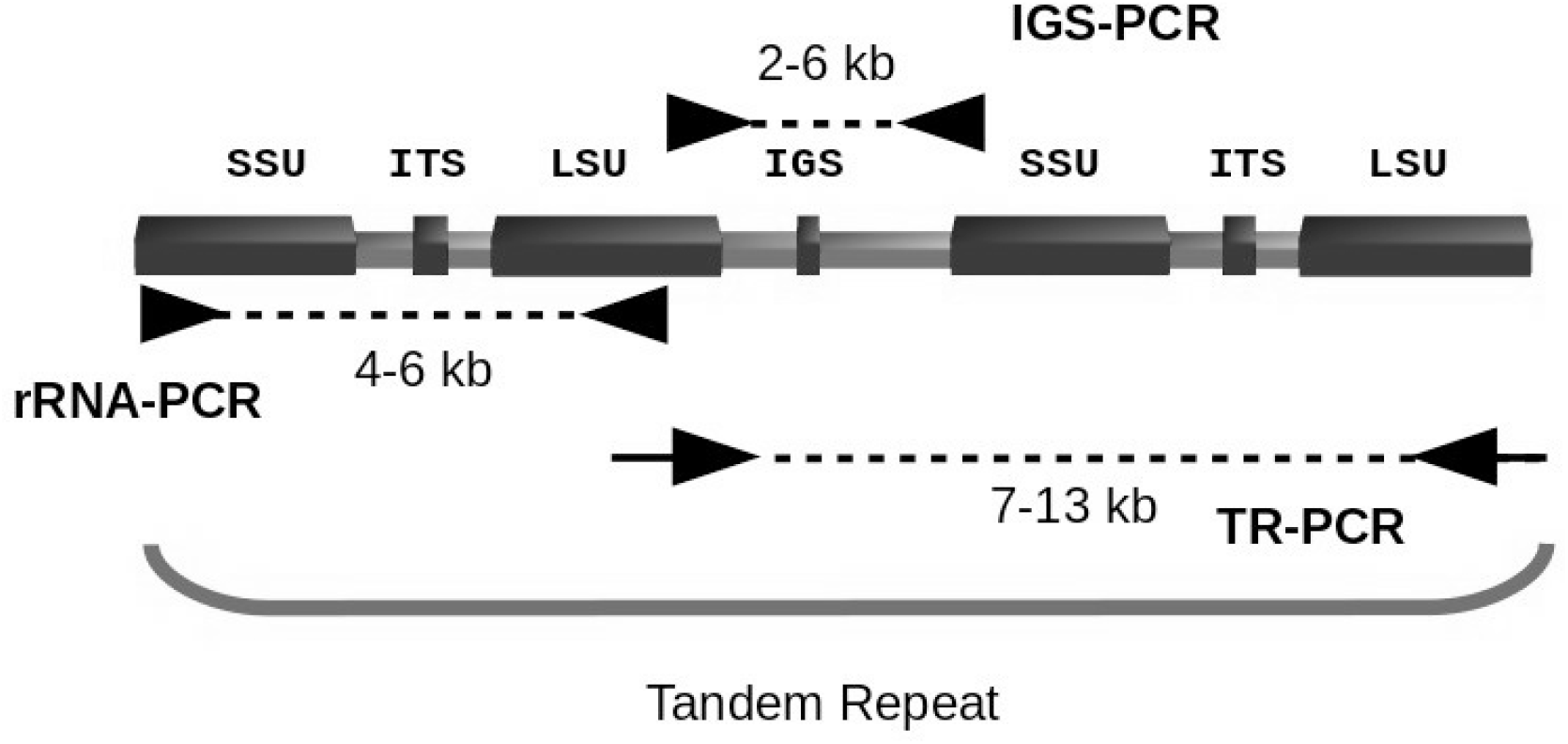
Schematic representation of the fungal ribosomal tandem repeat and our primer binding sites with two copies of the ribosomal operon. The three employed amplicons (rDNA, IGS, and TR) illustrate the covered regions.

Note that the last four nucleotides of both Fun-rOP primers are pairing and that these four nucleotides resemble the position overlap in the template (CTGA) at the exact *Escherichia coli* reference position 1770-1773 of the LSU (SILVA LSU reference position). This allows a subsequent end-to-end assembly of the full ribosomal region (LSU-IGS-ETS-SSU-ITS1-5.8S-ITS2-LSU) extracted from the ribosomal tandem repeat. All primer pairs were barcoded following the dual indexing strategy of Illumina sequencing (Part#15044223Rev.B; Illumina, San Diego). That means that we introduced the forward barcode series S500 to the 5’ end of each forward primer and the N700 barcode series to the 5’ end of the reverse primer, which allows the simultaneous sequencing of more than 100 samples, at least in theory. After each barcode we added one or two extra nucleotides as a precaution against nuclease activity. Between barcode and primer nucleotides we added a two-nucleotide wide spacer (see Supplemental Material S1) that has a mandatory mismatch to the fungal kingdom at these two positions. These two mismatches were validated by using ARB and the respective SILVA reference databases (SSU and LSU). We did not test all samples with all amplicons, since our focus in this study lay on the herbarium samples (Table 1).

**Table 1.** Amplification overview for the sample types

| technology | herbarium<br>specimens<br>(n = 66) | Chytridiomycota<br>cells (WGA)<br>(n = 9) | *Nephridiophaga*<br>(host tissue)<br>(n = 2) |
| --- | --- | --- | --- |
| PacBio | rDNA, IGS, TR | n.t. | rDNA |
| Nanopore | TR, rDNA | rDNA | n.t. |

n.t. not tested; rDNA refers to the amplicon produced by NS1short/RCA95m; IGS refers to the amplicon produced by RCA95rc/NS1rc; TR refers to the amplicon produced by Fun-rOP-F/Fun-rOP-T. WGA (whole genome amplification).

### Long range PCRs

In general, we applied the PrimeStar GLX polymerase (Takara) for all primer systems. For the IGS stretch (IGS PCR) and the full ribosomal region tandem repeat PCR (TR PCR), we employed only the PrimeStar GLX (Takara). The PCR was performed in 40 μl reactions with 1.5 μl enzyme, 12 pmol of each barcoded primer (a unique combination for each sample), 1 mM dNTP’s, and 1 μl of template (with a concentration of approximately 1-40 ng/μl). For all primer pairs we ran an initial denaturation of 1 min at 98°C, then 36 cycles at 98 °C for 10 sec, 55 °C for 15 sec, and 68 °C for 2.5 min. We increased the elongation step of the TR PCR from 2.5 to 4 minutes. For the Chytridiomycota samples we exchanged the PrimeStar polymerase for Herculase II (Agilent Technologies) for the rDNA PCR. The reason for this is that we worked on these samples in a second laboratory, in which the Herculase II was the established polymerase. We ran a two step protocol for Herculase II. An inital PCR with native (non-barcoded) NS1short/RCA95m primers and 3% BSA (molecular grade, Carl-Roth) as additive with 5 min at 95°C, then 35 cycles at 95 °C for 30 sec, 55 °C for 30 sec, and 68° C for 4 min. The PCR product was then used as template in a second PCR with 10 cycles but otherwise identical conditions, exchanging the native primers with barcoded primers.

### Library preparation and sequencing

The PCR products were purified with either 0.8 (v/v) of AMPure beads (Beckmann) or with PCR purification plates (Qiagen) following the respective manufacturer’s recommendations. After that, the purified PCR products were quantified using Nanodrop 2000 (Thermo Scientific) and pooled in an approximately equimolar way. This final pool was purified anew with AMPure beads using 0.4 (v/v) of beads and eluted in a 50-100 μl molecular grade water. The concentration of the amplicon pool was quantified with a Qbit instrument (Invitrogen). Approximately 2-4 μg were sent for sequencing with PacBio RSII (Pacific Biosciences) at the Swedish SciLife Lab in Uppsala, Sweden. Another batch of 800 ng was used for Oxford Nanopore library preparation following the manufacturer’s protocol and recommendations for D2 sequencing (LSK-208; Oxford Nanopore Technologies; discontinued as of May 2017, with the R9.4 chemistry) or alternatively 1D^2^ sequencing (LSK-308; with the most recent R9.5 chemistry as of May 2017). In brief, both protocols consist of end-repair, adapter ligation, and purification steps that take approximately two hours in the laboratory. Sequencing took place locally on a MinION instrument (Oxford Nanopore Technologies) operated with FLO-107 flowcells. We aimed for more than 2,000 sequences per sample and stopped the sequencing as soon as we achieved this goal, which took 2-8 hours depending on the pool size and amplicon length.

### Sequence data processing

The first step after obtaining the Nanopore data is the 2D basecalling, which was done with Albacore (v2.4; Oxford Nanopore Technologies). We observed that for a successful calculation of the required sequencing depth, usually 20% of the sequences are retained after this step as good quality 2D reads, which is one of the limitations of the 1D^2^ chemistry. Not all reads are complementary reads and currently the basecaller can only basecall ~50% as complementary (i.e. paired reads), while the other 50% remains unpaired. Unpaired reads have a higher error rate than paired reads and were therefore discarded in this study. These former as complementary identified sequences pair resulting in 25% of the initial reads. Finally, ~5% did not pass the quality filtering step, so that as a rule of thumb, a total of 20% of the initial reads remain as high quality paired reads for generating the consensus reference sequences.

For the PacBio data, we only work with the “reads of insert” (ROS) data in the next steps of the data processing. After these initial steps, all sequences from both sequencing platforms are processed in the same way. An initial quality filtering step (USEARCH v8.1; Edgar 2010) was performed. The maximum allowed error rate was set to 0.02 for PacBio sequences. After testing this quality filtering for the Nanopore data (which come pre-filtered at an error rate of 0.08), ranging from 0.04-0.08, however, we came to the conclusion that quality filtering had no beneficial effect on the final consensus quality (Supplemental Material S2). We thus removed it from the pipeline. Additionally, we filtered the sequences by length using biopython (v1.65; Cock et al. 2009) to exclude too short and too long sequences as detected in the histograms. This helped to increase the quality of the later alignment. Then all quality filtered and trimmed sequences were demultiplexed as FASTA files into individual samples according to their combined barcodes (Flexbar, v2.5, Dodt et al. 2012). Barcodes and adapters were removed in this step. All sequences from each individual sample were subsequently aligned using MAFFT (v7, Katoh & Standley 2013) using the autoalignment option. The aligned sequences were clustered in Mothur (v1.39, Schloss et al. 2009) using the Opticlust algorithm, and the consensus sequences for each operational taxonomic unit were built using a custom-made Perl script (Consension) available at http://microbiology.se/software/consension/. The optimal OTU clustering threshold for Nanopore data was determined to be 0.07 for shorter amplicons (rDNA and IGS PCR) and 0.08 for the long TR PCR (Supplemental Material S3). To counter spurious OTUs we determined a dynamic OTU size cut-off that is given to Consension, which was calculated as:

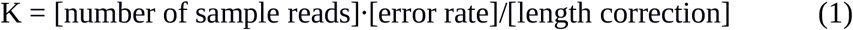

The length correction is an integer and defined as amplicon length [kb] divided by 5 kb of the expected amplicon length. K_min_ (the minimum number of sequences a OTU can hold) was set to 3 for PacBio and 5 for Nanopore sequences. The consensus sequences were finally compared by inspecting the alignment visually (Gouy et al. 2009) and by calculating sequence similarities with local BLAST searches (nucleotide BLAST, v2.2.28). Visualization of the BLAST-based TR results matching all other sequences (rDNA, IGS, and ITS) was done with BRIG (v 0.95, Alikhan et al. 2011). In the few cases where we obtained more than 1 OTU after consensus generation, we only used the most abundant OTU for subsequent similarity comparisons. Finally, we evaluated the effect of polishing the Nanopore consensus sequences by mapping the FASTQ files to them with Racon (https://github.com/isovic/racon).

## Results

All primer pairs worked successfully on our target herbarium fungal samples. The rDNA primers also worked with samples from the early diverging lineages of Chytridiomycota (whole genome amplified DNA of infected single algal cells) and host tissue infected with *Nephridiophaga.* This confirmed the fungal specificity and the broad spectrum of the primers, which should, based on *in silico* analysis, cover all fungal phyla with the few within-phyla exceptions mentioned above (see Primer design). We noted, however, that the whole ribosomal region (TR PCR) was amplified in only 50% of the tested herbarium specimens (29 of 58 samples), potentially due to DNA integrity issues (see Discussion section and Larsson & Jacobsson, 2004).

An example of a full comparison between Sanger, PacBio, and Nanopore-generated sequences and all applied primer pairs for one of our herbarium DNA samples can be seen in Figure 2 for specimen GB-0158876 *(Inocybe melanopus* EL263-16).

**Figure 2.**
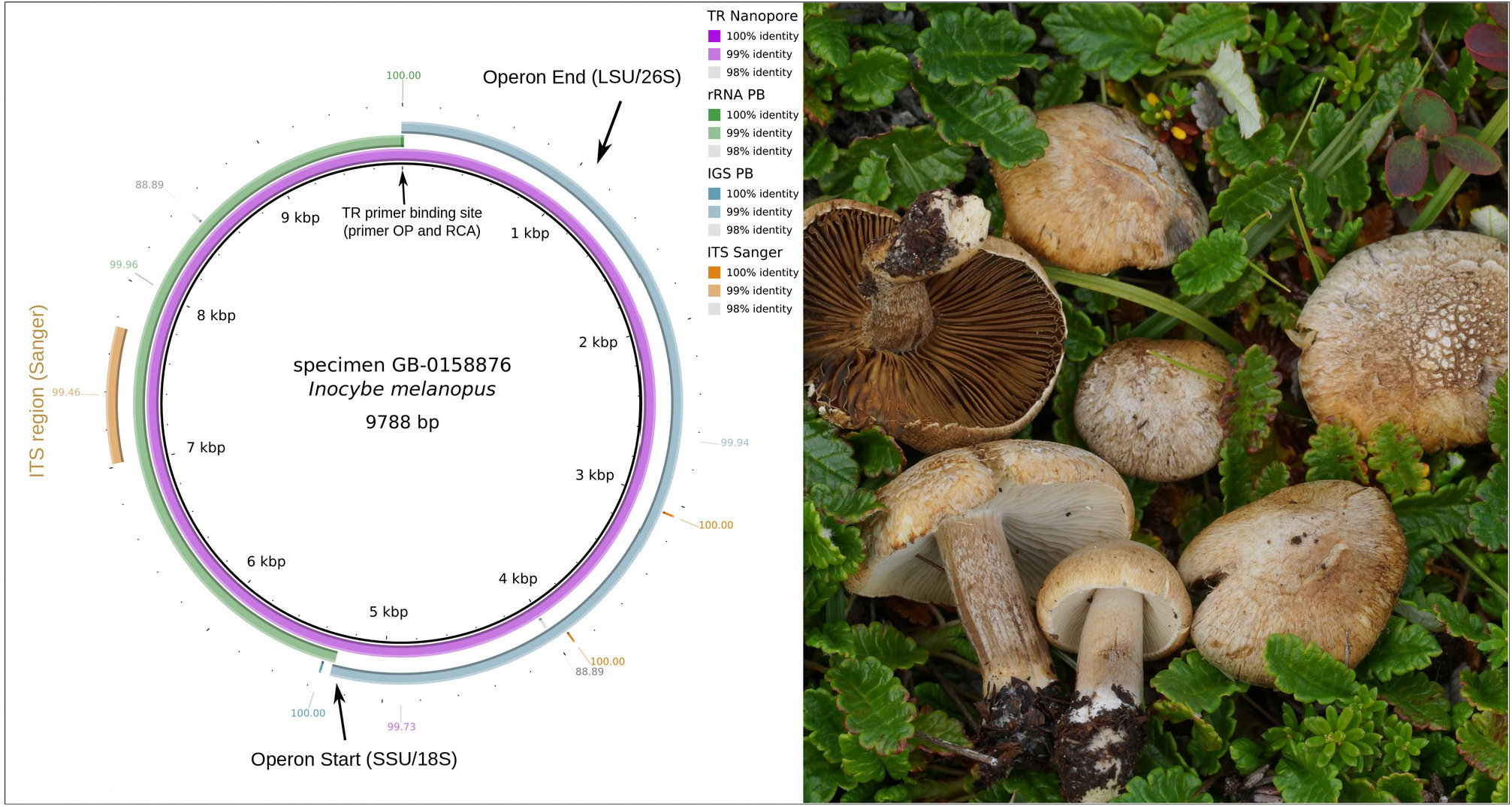
Left panel: graphic view on the BLAST comparison of the ribosomal regions generated with the three amplicons and the Sanger reference sequence of the ITS region. The TR amplicon generated by PacBio sequencing was set as reference. Similarities are displayed for each fragment, respectively. Right panel: photograph of in situ basidiomata of herbarium specimen GB-0158876 *(Inocybe melanopus).*

As expected, PacBio-derived sequences had a high accuracy and matched good quality Sanger sequences with 100% identity in 23 of 64 cases, while Nanopore sequences achieved this only in 3 of 41 cases. The discrepancy between PacBio and Sanger was in most cases related to mismatches in the distal ends of the ITS sequences, potentially reflecting quality issues of Sanger sequences (Figure 3). Similarly, Nanopore-derived sequences (1D^2^ chemistry) had on average only 0.15% mismatches to PacBio sequences, resulting in a consensus accuracy of 99.85%, identical to the median Sanger similarity to PacBio sequences (Table 2). In the alignment view, most of the Nanopore-based mismatches could be identified as indels in homopolymeric regions. The discontinued D2 chemistry in combination with the outdated base calling reached a consensus accuracy of 99.4%.

**Figure 3.**
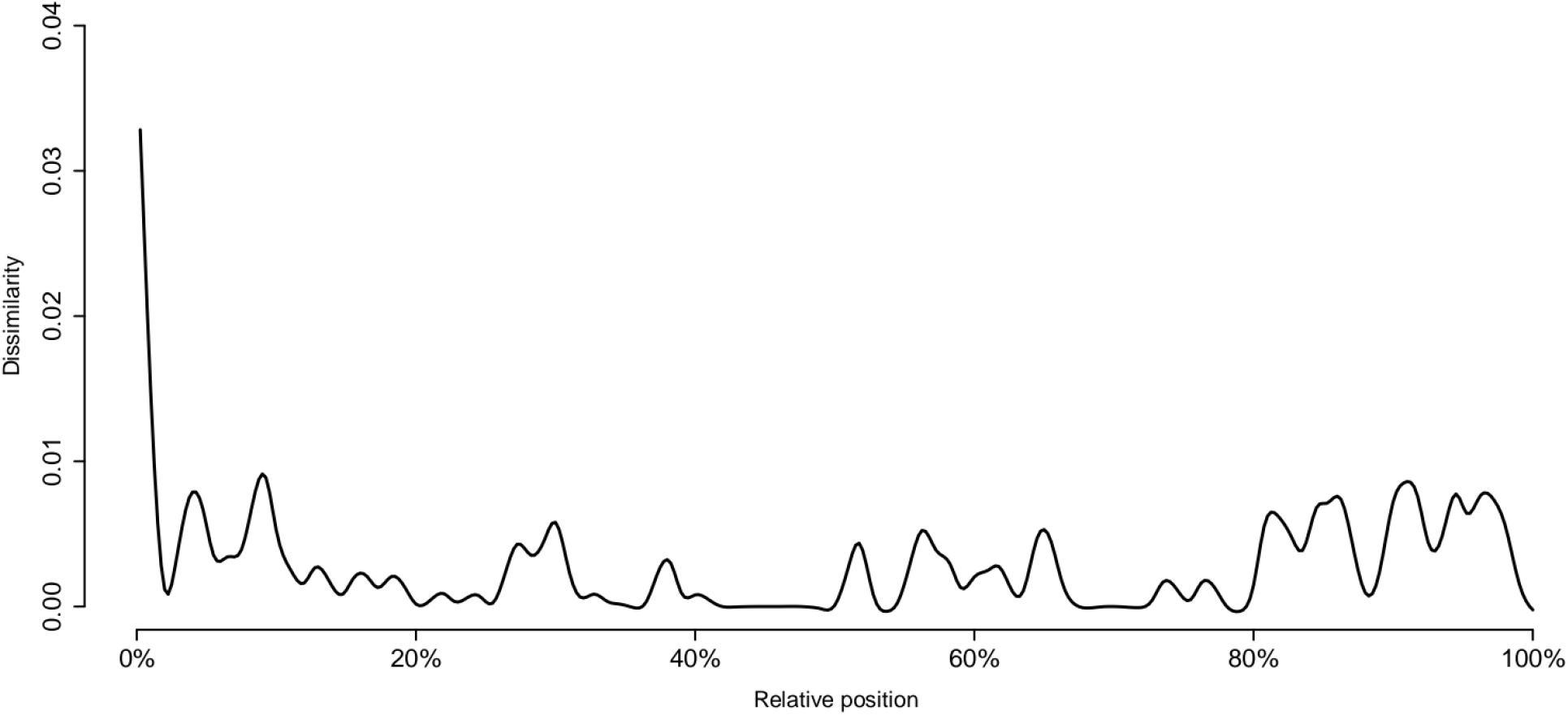
Quality graph of ITS Sanger sequences versus PacBio references, that show a small bias at the 5 prime end of the ITS fragment. The graph is based on 56 Sanger-PacBio sequence pairs.

**Table 2.**
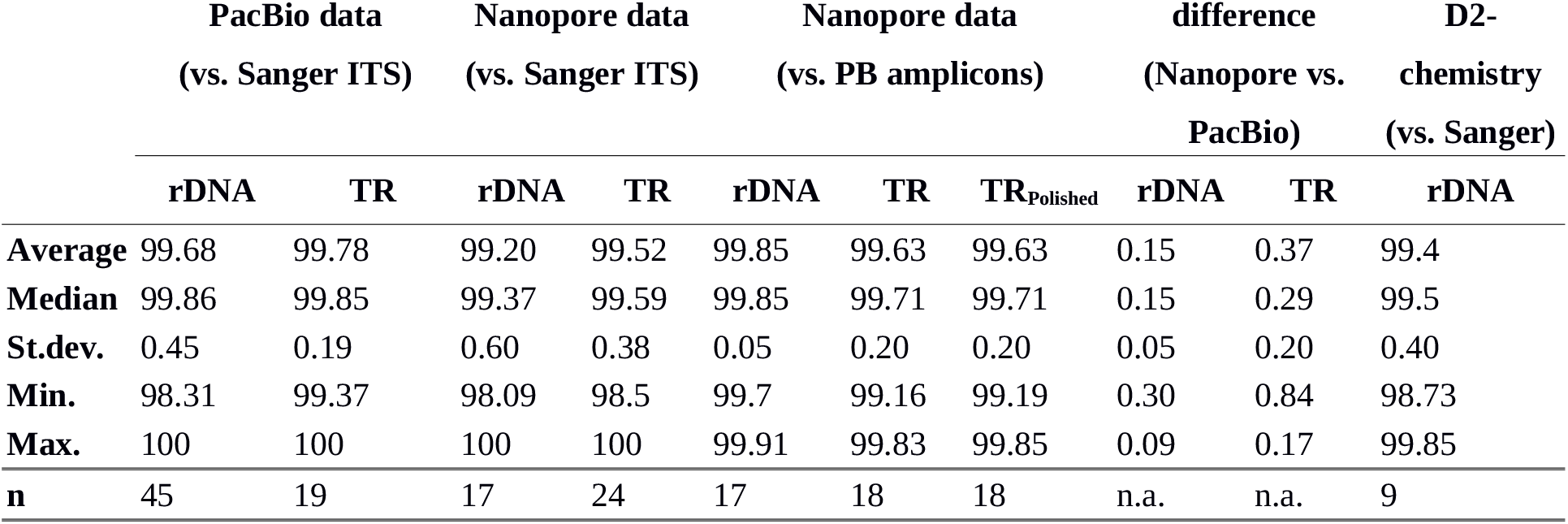
Similarities of Sequences compared to Sanger or PacBio sequences

|  | PacBio data<br>(vs. Sanger ITS) |  | Nanopore data<br>(vs. Sanger ITS) |  | Nanopore data<br>(vs. PB amplicons) |  |  | difference<br>(Nanopore vs.<br>PacBio) |  | D2-chemistry<br>(vs. Sanger) |
| --- | --- | --- | --- | --- | --- | --- | --- | --- | --- | --- |
|  | rDNA | TR | rDNA | TR | rDNA | TR | TR<sub>Polished</sub> | rDNA | TR | rDNA |
| **Average** | 99.68 | 99.78 | 99.20 | 99.52 | 99.85 | 99.63 | 99.63 | 0.15 | 0.37 | 99.4 |
| **Median** | 99.86 | 99.85 | 99.37 | 99.59 | 99.85 | 99.71 | 99.71 | 0.15 | 0.29 | 99.5 |
| **St.dev.** | 0.45 | 0.19 | 0.60 | 0.38 | 0.05 | 0.20 | 0.20 | 0.05 | 0.20 | 0.40 |
| **Min.** | 98.31 | 99.37 | 98.09 | 98.5 | 99.7 | 99.16 | 99.19 | 0.30 | 0.84 | 98.73 |
| **Max.** | 100 | 100 | 100 | 100 | 99.91 | 99.83 | 99.85 | 0.09 | 0.17 | 99.85 |
| **n** | 45 | 19 | 17 | 24 | 17 | 18 | 18 | n.a. | n.a. | 9 |

In a few samples (on average in 12% of the amplicons) we saw two or more distinct consensus sequences, both of which crossed the consensus threshold K. Often these lower abundance alternative consensus sequences differ significantly – by more than 100 bases - in length, and it is likely that these represent genomic variants of the ribosomal regions. Currently, we disregard these sequences, but they may be worth a second look in future studies, as they may have implications for the outcome of phylogenetic analyses. We found that a lower clustering threshold (0.08 or below) is important in keeping special cases of these variants, i.e. identical sequences with one large intron, separate (Supplemental Material 3).

## Discussion

The distribution of ribosomal marker sequences across distinct databases is not only a mycological problem but one that pertains to most DNA-based studies targeting bacteria, algae, and protists (e.g. De Vargas et al. 2015; Wurzbacher et al. 2017). We are currently missing a lot of reference data in the databases used on a regular basis. In a time where biodiversity screening by high-throughput sequencing methods is becoming routine, these deficits in taxonomic and marker-related sampling of genetic material are developing into severe problems. Any strategy that may help closing these gaps in the future will be extremely valuable. The genetic markers of the ribosomal operon differ from each other, and across the fungal tree of life, in length and level of conservation. As a consequence, different markers have been used to address research questions in different parts of the fungal tree of life. The incompleteness and fragmentation of extant ribosomal data is clearly problematic, and our sampling of fungal ribosomal DNA sequences should be augmented with full-length reference sequences that span several regions suitable for everything from conservative taxonomic classification to intraspecies assignment. The primers we present here extend the currently longest sequenced ribosomal fragments (Karst et al. 2018, Tedersoo et al. 2018), and enable generation of data in an easy and straightforward way for the whole fungal ribosomal tandem repeat region, solving the problem of non-overlapping sets of ribosomal markers.

The primer pairs we introduce with the present paper worked not only for the Basidiomycota species examined, but also for Chytridiomycota and the distant genus *Nephridiophaga.* Indeed, according to the *in silico* primer design and evaluation, they should be suitable for the greater majority of fungal species, with the exception of long-branching Zygomycota lineages, for which primer adaptations may be required. The primer RCA95m and the OP primers are fairly fungus-specific and help to amplify fungi from non-fungal DNA (e.g. our cockroach host tissue, see also Heeger et al. 2018). The decreased success rate that we see for the TR amplification in comparison to the rDNA and IGS PCR is probably linked to the DNA integrity in the sense that DNA is known to degrade (fragment) over time in herbarium settings (Larsson & Jacobsson, 2004). The TR-PCR approach requires long genomic fragments with intact ribosomal tandem repeats. Here, we used herbarium samples, which are usually moderately to fairly fragmented depending on age of collection and storage conditions, so the degree of fragmentation may have hindered the amplification of the TR fragment, while the shorter rDNA and IGS PCRs still worked. Although the accuracy of TR fragments (99.63 %) is slightly lower, it is still in the range of Sanger sequencing. A polishing step was not successful in increasing the overall quality (Table 2). The reason could lie in the underlying alignment algorithm, which may not be optimized for long fragments with high individual error-rates. In summary, the concerted sequencing of rDNA and IGS is currently a robust way in obtaining the complete ribosomal fragment in terms of sequence quality and in cases of lower template DNA integrity.

Long-read sequencing offers the possibility to generate reference data for fungi, potentially other eukaryotes, and bacteria at high read quality. Assuming that our PacBio data is almost perfect (99.99% accuracy, Travers et al. 2010), the average Nanopore consensus quality of 99.85% is already as good as the average Sanger quality of 99.78% (Nilsson et al. 2017) or for our data 99.73% (average of TR and rDNA result, Table 2). We argue, therefore, that the use of Nanopore sequencing is justified. In particular, Nanopore sequencing offers a cheap method to generate full-length ribosomal data, independently of amplicon length, at excellent quality and does not require additional clean-up steps, as may be necessary for PacBio (See Supplemental Material S4 for a direct comparison between PacBio and Nanopore in terms of produced fragment lengths). We anticipate that Nanopore sequencing will prove to be a valuable tool for small laboratories, culture collections, herbaria, field work, and single cell workflows for the generation of high-throughput reference data, potentially also for mixed environmental samples (Karst et al. 2018, Calus et al. 2018). The sequencing can be done in-house within a couple of days, significantly speeding up the generation of reference data and rendering it suitable for high-throughput solutions. Importantly, this approach does not rely on sending DNA for sequencing at large-scale facilities but is, rather, amenable to analysis on a modern desktop computer.

Similar to mock communities in environmental samples (Heeger et al. 2018), we consider it as mandatory for Nanopore generated data to spike in control DNA, if no partial reference data is available, or do complementary ITS Sanger/PacBio sequencing as an internal standard control. The generated data should be deposited as carefully as in the case of Sanger sequences to avoid errors derived from the experimental procedure (cf. Nilsson et al. 2017). The sequencing error rate, determined through the use of known high-quality reference data, should always be included in the submission, included either in the sequence header or as additional data.

In conclusion, we hope that this manuscript lays out the first steps for a new way of generating full-length reference data for fungi. This will enable mycologist to comprehensively fill up the taxonomic and marker-related gaps in the fungal databases in a straightforward and cost-efficient way. We found the adaptation to high-throughput data generation to be surprisingly easy, and to require only an initial investment in barcoded primers, as well as good sample and data management. Our approach does place some demands on availability of bioinformatics expertise, testifying to the multidisciplinary nature of contemporary mycology.

## Acknowledgements

We would like to thank Renate Radek for her help with providing material of *Nephridiophaga;* Keilor Rojas-Jimenez for whole genome amplifications; and Magnus Alm Rosenblad, Michael M. Monaghan, Elizabeth C. Bourne, and Felix Heeger for joint discussions on the implementation of long-read sequencing. The authors would like to acknowledge support from Science for Life Laboratory, the National Genomics Infrastructure, NGI, and Uppmax for providing assistance in massive parallel sequencing and computational infrastructure. CW and RHN gratefully acknowledge financial support from Stiftelsen Olle Engkvist Byggmästare, Stiftelsen Lars Hiertas Minne, Kapten Carl Stenholms Donationsfond, and Birgit och Birger Wålhströms Minnesfond. SVdW was supported by a IGB postdoc fellowship and the German Science Foundation (DFG).

## Footnotes

1 We noticed that there are a few cases in the reference data where this 5’ prolongation “CA” is replaced by “TC” in several not necessarily related fungal species. Closer inspection of these cases by BLASTing and aligning to the SILVA SSU reference database (v. 128) showed that this was due to incomplete trimming of the sporadically used primer PNS1 (Hibbett DS. 1996. Phylogenetic evidence for horizontal transmission of Group I introns in the nuclear ribosomal DNA of mushroom-forming fungi. Mol Biol Evol 13:903-917.), which produces severe 5’ mismatches to the fungal backbone.

